# An information theoretic approach to detecting spatially varying genes

**DOI:** 10.1101/2022.11.02.514777

**Authors:** Daniel C. Jones, Patrick Danaher, Youngmi Kim, Joseph M. Beechem, Raphael Gottardo, Evan W. Newell

## Abstract

Identifying genes with spatially coherent expression patterns is a key task in spatial transcriptomics. We adopt an information theoretic perspective on this problem by equating the degree of spatial coherence with the mutual information between nearby expression measurements. To avoid the notoriously difficult problem of computing mutual information, we use modern methods of approximation, in a method we call maximization of spatial information (Maxspin). As well as being highly scalable, we demonstrate improved accuracy across several spatial transcriptomics platforms and a variety of simulations when compared to both existing specialized methods and traditional spatial statistics methods. We use the method to analyze a renal cell carcinoma sample profiled using CosMx Spatial Molecular Imaging, revealing previously undescribed gene expression patterns.

---

The last several years have seen a slew of advancements in spatially resolved transcriptomics [Marx, 2021, Moses and Pachter, 2022], proteomics [Lundberg and Borner, 2019], and genomics [Zhao et al., 2022]. These methods create an opportunity to develop a deep understanding of the spatial organization of tissues at the cellular level. When the expression of a large number of genes is measured, a key consideration is identifying which genes show signs of structured, nonrandom spatial organization. Analogously to identifying differentially expressed genes, these methods aim to identify spatially varying genes (SVGs).

Similar questions have been examined in geostatistics, ecology, and demography for decades, so naturally, existing spatial statistics methods have been put to use. Traditional measures of spatial autocorrelation, the most common of which are Moran’s I and Geary’s C [Ripley, 2005], have numerous general purpose implementations, but have also been included in spatial expression analysis toolkits like Squidpy [Palla et al., 2022] and MERINGUE [Miller et al., 2021].

A number of methods have built more elaborate, special-purpose models based on Gaussian processes with covariance functions applied over spatial coordinates (“kriging” in geostatistics parlance), which have long been a standard tool in probabilistic modeling of spatial data. SpatialDE [Svensson et al., 2018] tests for SVGs by separately fitting a Gaussian process model and a regression model excluding spatial covariance, and comparing the two using a χ^2^ test. SPARK [Sun et al., 2020] elaborates on this primarily by using Poisson likelihood, which is a more natural model of count data, and by testing models with ten different covariance functions to account for various potential patterns of spatial coherent expression. GPcounts [BinTayyash et al., 2021] models counts using a negative binomial distribution, and tests for spatial coherence using a likelihood ratio test on the marginal likelihoods of models with and without the Gaussian process prior.

Gaussian process models have many appealing properties, but also one major shortcoming: exact inference is O (n^3^), where here n is the number of cells or spots. This quickly becomes problematic once n is larger than a few thousand. Given the recent rapid progress in spatial assays, cubic time inference marks a method for rapid obsoletion. A recent method, nnSVG [Weber et al., 2022], overcomes this problem by leveraging a modern approximate inference scheme for Gaussian process models. SPARK-X [Zhu et al., 2021] instead bypasses the issue by using an entirely different model than SPARK, a null-hypothesis test on the Frobenious inner product of the spatial coordinate covariance matrix and one computed over a gene’s expression. When these are highly-nonindependent, the inner product test statistic will be large, yielding smaller p-values. To consider various patterns of spatial coherence, it runs this test on different transformations of the coordinates.

Apart from the Gaussian process methods, trendsceek [Edsgärd et al., 2018] uses a mark-segregation test, which tests the statistical independence of the distance between two cells, and a gene’s expression in both. If conditioning on the distance does not alter the distribution over expression pairs, there is no detectable spatial coherence. Independence is tested using summary statistics, and p-values are computed by shuffling expression values across spatial locations to compute an empirical null distribution. Sepal [Anderson and Lundeberg, 2021] constructs a model simulating continuous time diffusion of expression across the spatial domain and measures spatial coherence by the time required for the process to reach a homogeneous state.

Giotto [Dries et al., 2021], in addition to interfacing with SpatialDE, trendsceek, and SPARK, implements its own SVG test called binary spatial extraction (BinSpect) which binarizes expression values using either K-means clustering (BinSpect-kmeans) or threshold ranking (BinSpect-rank) and runs a Fisher’s exact test using a contingency table of values from neighboring cells. SpaGene [Liu et al., 2022] also heuristically binarizes expression data, but instead considers the degree distribution for a subgraph of the k nearest neighbor graph consisting of just high expression cells, operating under the principle that if high expression cells colocate we would expect an increase in higher degree nodes in this graph when compared to a shuffled graph.

Existing methods often make either strong distribution assumptions, discard information through binarization, or are highly tuned in an ad-hoc manner to specific types of patterns. In the work presented here, we develop a entirely new approach to measuring spatial coherence, adopting an information theoretic perspective and building on recently advancements in approximating mutual information. Like trendsceek our goal is to quantify the degree of statistical dependence between a pair expression expression values and their spatial vicinity, but rather than running null hypothesis tests on summary statistics, we take a more direct route by estimating the mutual information between the two. Spatial coherence can then be neatly defined as how much information gene expression values in the same neighborhood share. In a spatially incoherent setting, mutual information is zero; nearby expression values share no more information than randomly selected values. When expression is organized spatially, expression pairs become increasingly predictable based on whether they are nearby or not, and mutual information will be high.

Mutual information is appealing on theoretical grounds, but its application is often fraught. Computing mutual information directly necessitates a model of the joint probability, and in the case of continuous domains, computing a multidimensional integral. Recently, mutual information has gained renewed interest in the context of deep learning [Tschannen et al., 2019]. Belghazi et al. [2018] showed, in a method they call mutual information neural estimation (MINE), how mutual information between to variables X, Y can be estimated by training a neural network classifier to distinguish pairs of observations (de facto draws from the joint distribution P(X, Y)), from pairs drawn from the marginal distributions P(X), P(Y), which can be formed by shuffling observations. Intuitively, the better the neural network is able to distinguish observed pairs from shuffled pairs, the higher the mutual information. This intuition is formalized by showing that the objective function used to train the neural network forms a lower bound on the mutual information.

In our method, Maxspin (a portmanteau of “maximization of spatial information”), we adopt the MINE approach to compute lower bounds on the mutual information between pairs of nearby gene expression values. On a genewise basis, we optimize a simple classifier to distinguish pairs of expression value pairs chosen uniformly at random across spatial locations, from pairs chosen from nearby locations according to a short random walk on the spatial neighborhood graph (Figure 1a). Spatial coherence is scored as the degree to which a classifier can recognize uniformly sampled pairs from spatially proximate pairs, with an objective function that is a lower bound on the Kullback-Leibler divergence between the two distributions, representing a the spatial auto mutual information, which we will refer to as simply “spatial information” here. This principle can be trivially generalized to consider the spatial information between pairs of genes, quantifying the degree of spatial coexpression.

**Figure 1:**
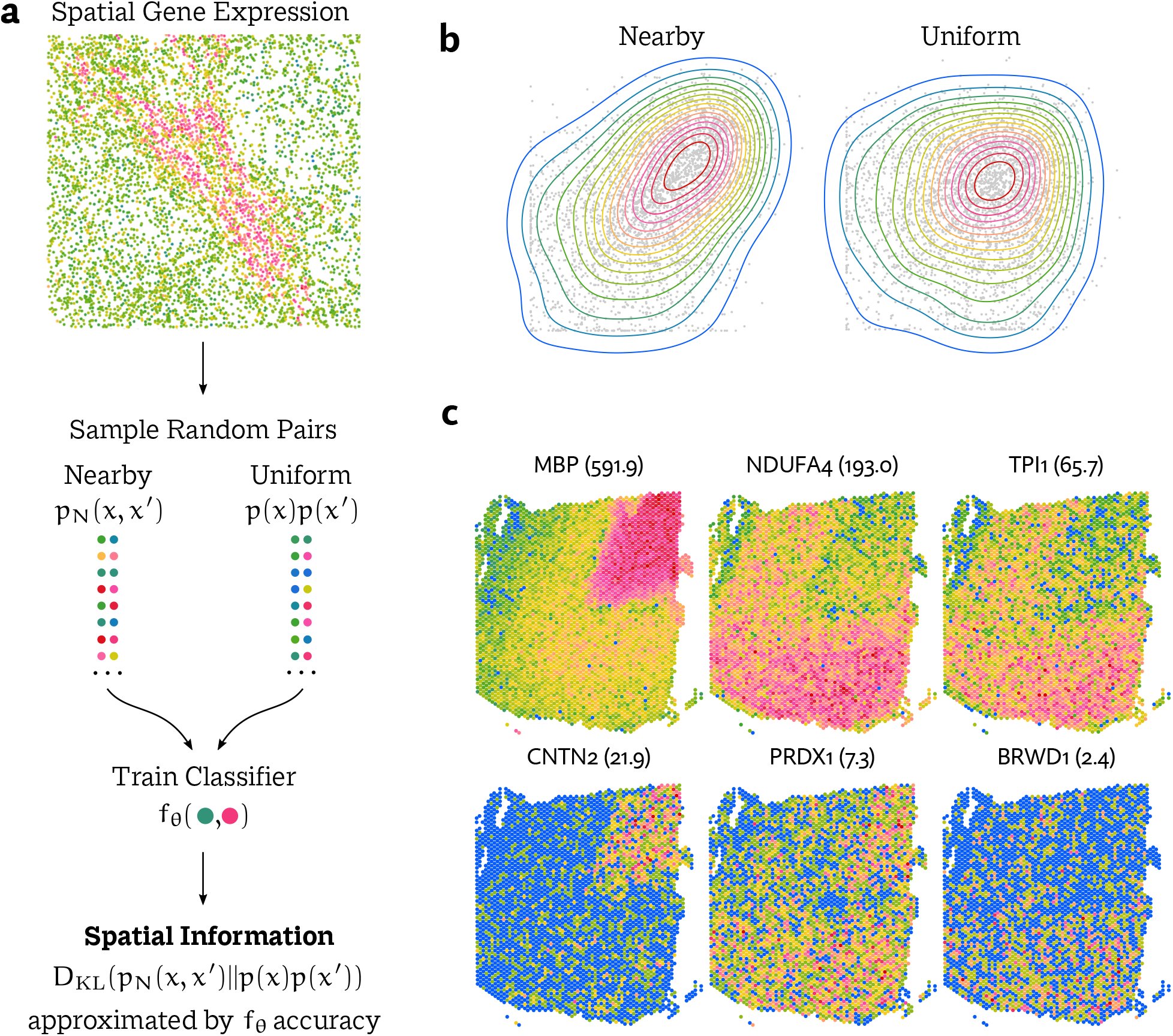
**a** The maximization of spatial information (Maxspin) algorithm proposed here works by training a simple classifier to distinguish between expression pairs sampled from nearby locations and those chosen uniformly at random. **b** If a gene displays spatial organization there is a distributional shift between the two sets of sampled cell pairs. The classifier accuracy acts as an approximation of the Kullback-Leibler divergence between these distributions, which can be interpreted as spatial information. **c** The spatial information score computed effectively quantifies the degree of spatial coherence for each gene evaluated, here shown in a Visium mouse brain data set.

## Results

To evaluate the performance of Maxspin and existing methods of detecting spatially varying genes, we used three publicly available data sets generated using different spatial transcriptomics technologies: 10X Visium, NanoString CosMx SMI, and MERFISH. We additionally developed a new simulation methodology and examined a large variety of simulations, in addition to more simplistic simulations originally used to evaluate SPARK-X.

In each of the three data sets we have some reasonable annotation of spatially organized cell type or region. As a proxy for ground truth, we called differential expression between pairs of annotated cell groups, producing a list of spatially varying genes. This list omits some SVGs with spatial expression patterns not conforming to the annotated regions, but we expect it to contain the large majority of the most obvious examples, and omissions should equally disadvantage each method, not a priori biasing the benchmark towards any particular method. Differential expression was called using pairwise Wilcoxon signed-rank tests, using a q-value cutoff of 0.01 after applying the Benjamini-Hochberg correction.

Because of the wide variety of settings, we adopt Moran’s I as a baseline method and consider the degree to which a method improves on (or falls below) its performance, as measured by the area under the precision-recall curve (PR-AUC). Moran’s I is simple, easy to interpret and implement, and has been a mainstay of spatial statistics for seventy years. Thus it’s a meaningful hurdle that any specialty method for detecting spatially varying genes should aim to surpass.

GPcounts was excluded from these benchmarks. Though it has a test of spatial coherence, it was designed for very small numbers of cells and fails to scale to any of the datasets evaluated here. We were also unable to run trendsceek as the software is unmaintained to has been rendered unusable by library incompatibilities.

### Visium

We used human prefrontal cortex data from Maynard et al. [2021], generated on the 10X Visium platform. The expression of 21,151 genes was profiled in 12 samples. Visium slides consist of a grid of 4992 spots laid out in a hexagonal pattern. In this data set between 3460 and 4789 spots were covered by the tissue samples. Compared to single-cell methods, Visium spots are typically much more deeply sequenced, but are lower resolution, containing several cells in most cases. To determine a ground truth set of SVGs, we used the authors’ annotation of cortical layers, calling differential expression between all pairs of adjacent layers, resulting in 7775 spatially varying genes on average.

Every method considered was able to scale to the Visium data by virtue of having a strict maximum number of spots. In future generations of the technology with more spots, this will become more problematic for SPARK and SpatialDE, for which this dataset pushes the limit on tractability. Overall, we demonstrate consistently improved accuracy using Maxspin (Figure 2a). Following Maxspin, the two Gaussian process methods SpatialDE and nnSVG perform well, while SPARK-X was inconsistent, showing high accuracy some of the 12 samples, but greatly diminished for others. SPARK and BinSpect offer only very small improvement’s over Moran’s I, while Geary’s C and SpaGene tend to underperform it. Sepal requires spatial data on a regular grid, so we were only able to apply it to this dataset, however its performance was exceedingly poor so was omitted here for clarity.

**Figure 2:**
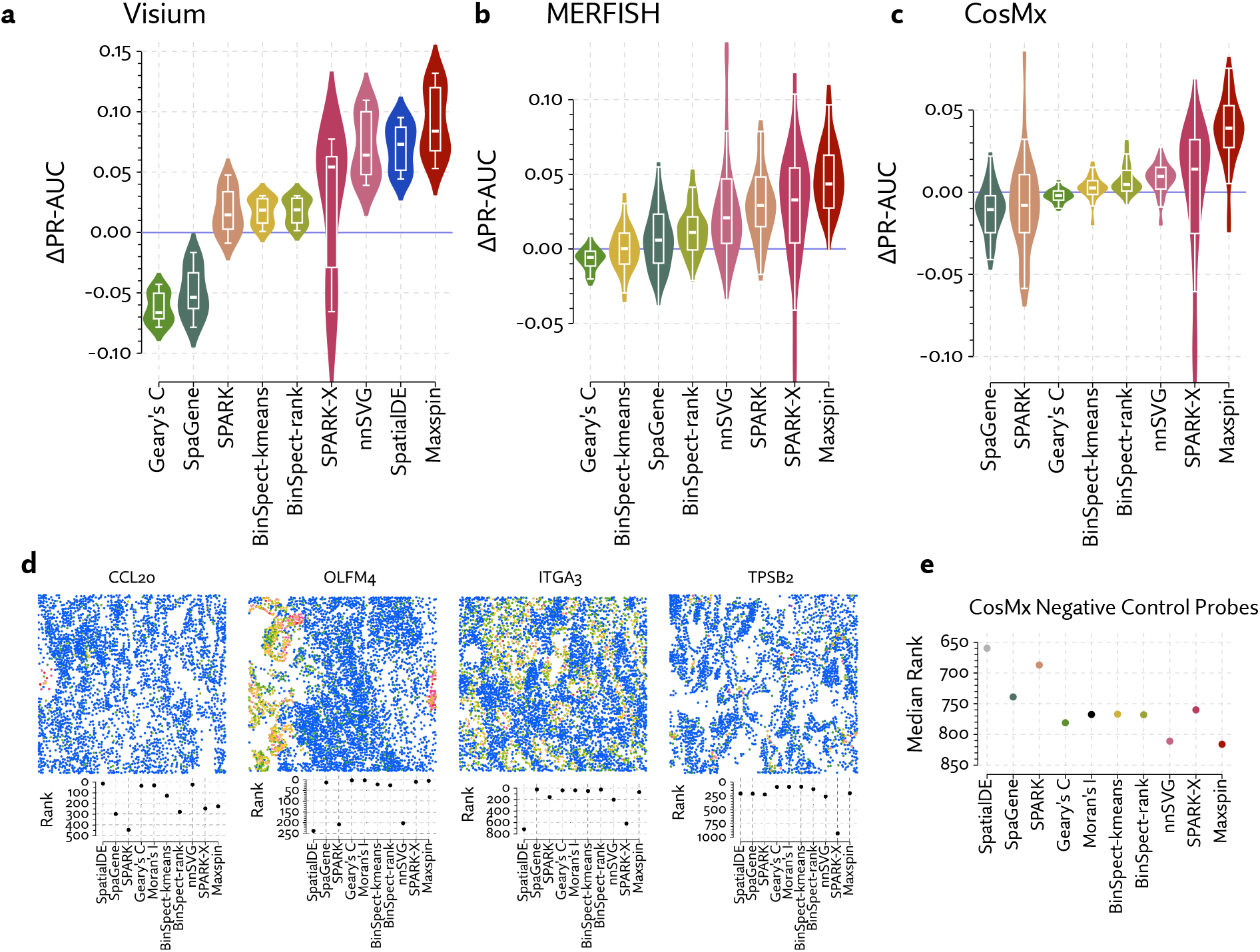
Methods for detecting spatially varying genes were evaluated by calling differential expression between manually annotated regions, and treating this list as ground truth. The area under the precision-recall curve (PR-AUC) was computed for each test, and the value achieved by Moran’s I was subtracted, producing ΔPR-AUC, representing the improvement over this baseline method. Methods were evaluated in multiple data sets generated using **a** Visium, **b** CosMx, **c** and MERFISH. Results for SpatialDE in the MERFISH and CosMx benchmark were exceedingly low, so excluded from these plots. **d** Selected examples of disagreements between methods in the CosMx dataset. e Median ranks (out of 960 genes) of negative control probes in the CosMx dataset, which should not be significantly expressed nor spatially coherent.

In Figure 1c, a span of information scores computed by Maxspin is shown for one sample in this dataset.

### MERFISH

Mouse primary motor cortex MERFISH data was taken from Zhang et al. [2021]. The data consists of panel of 252 genes measured in 12 experiments, each consisting of a number of coronal slices (64 in total), with a total of 280,186 cells. We annotated cortical layers by reproducing the procedure described by the authors, discarding one of the experiments where we could not reliably annotate the L6b layer. For each experiment, we performed pairwise differential expression between all adjacent pairs of cortical layers, taking the union to produce a list of spatially varying genes that were treated as ground truth for these benchmarks. Because this dataset consists of a smaller gene set, which is selected to be enriched for genes varying across layers, most of the genes show differential expression between at least one pair of layers. To create a somewhat more subtle test, instead of considering whole slices, we instead benchmarked the 180 pairs of adjacent layers from each slice. The ground truth set of spatially varying genes consisted of an average of 110 genes.

Maxspin has the highest median improvement over Moran’s I. SPARK-X is not far behind in terms of median ΔPR-AUC, but with a considerably higher proportion of low scoring examples. The Gaussian process methods SPARK and nnSVG also improve on Moran’s I, though to a lesser extent. The remaining methods are comparable to Moran’s I. Departing from the Visium results, SpatialDE displayed exceedingly poor performance, with a median ΔPR-AUC of −0.139. The cause of this appears to be a tendency to assign p-values of 0 to a large proportion of the genes, thus lacking any ability to order these meaningfully.

Supplemental Figure 4 shows examples of genes with a range of spatial information scores, the highest scoring genes correspond to starkly elevated expression in particular layers, and the smallest scores to a much subtler layer-wise enrichment.

**Figure 3:**
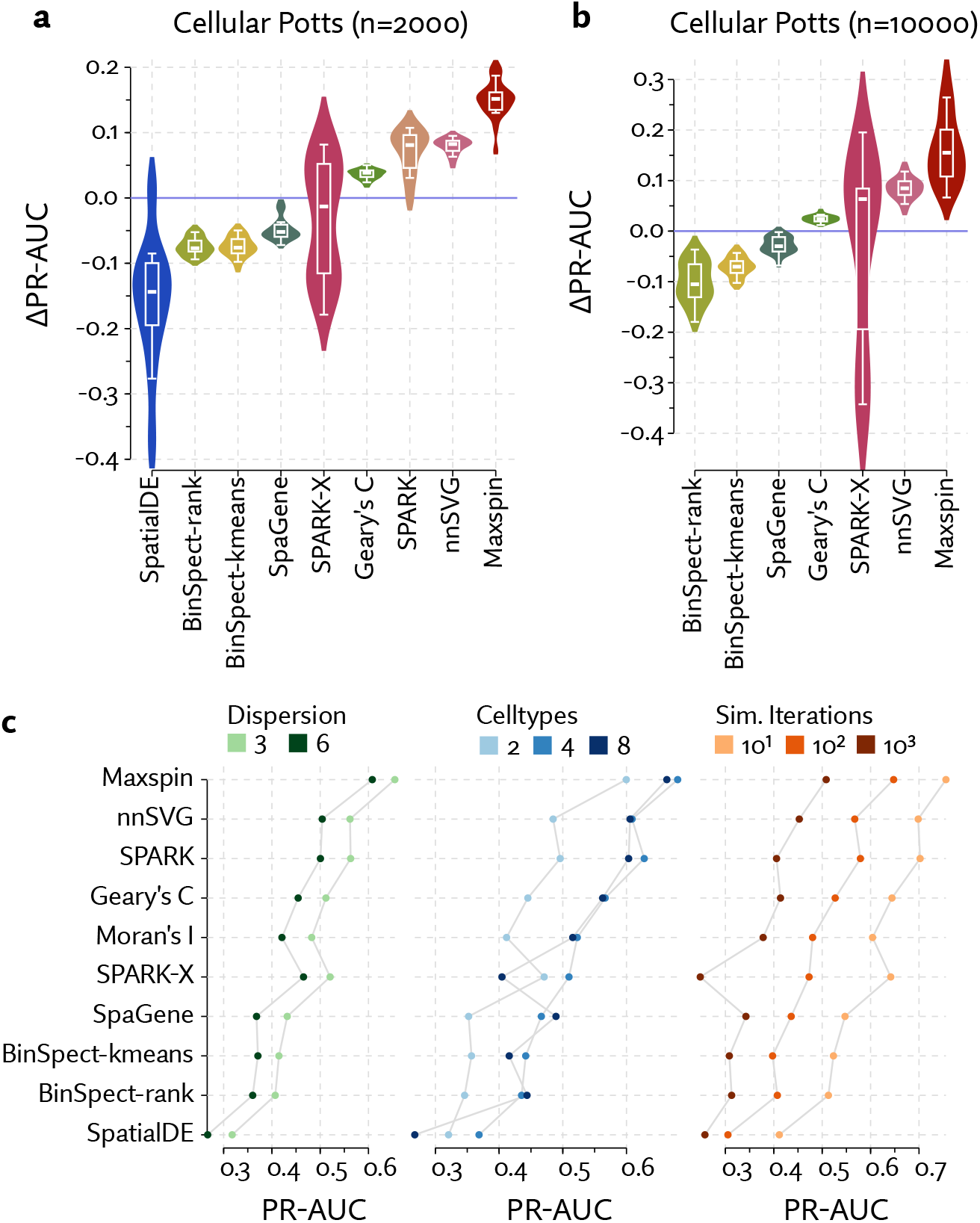
Spatial transcriptomics data was simulated using a cellular Potts model for **a** 2000 and **b** 10000 cells, with expression drawn from a negative binomial distribution that is perturbed across cell types for the ground truth spatially varying genes. Methods were benchmarked by computing the area under their precision-recall curves and subtracting the value achieved by Moran’s I. **c** Performance is further broken down by various parameterizations of the simulation. Across the board, performance increases with lower dispersion, fewer simulation iterations (resulting in simpler spatial distributions), with number of cell types playing a more ambiguous role.

**Figure 4:**
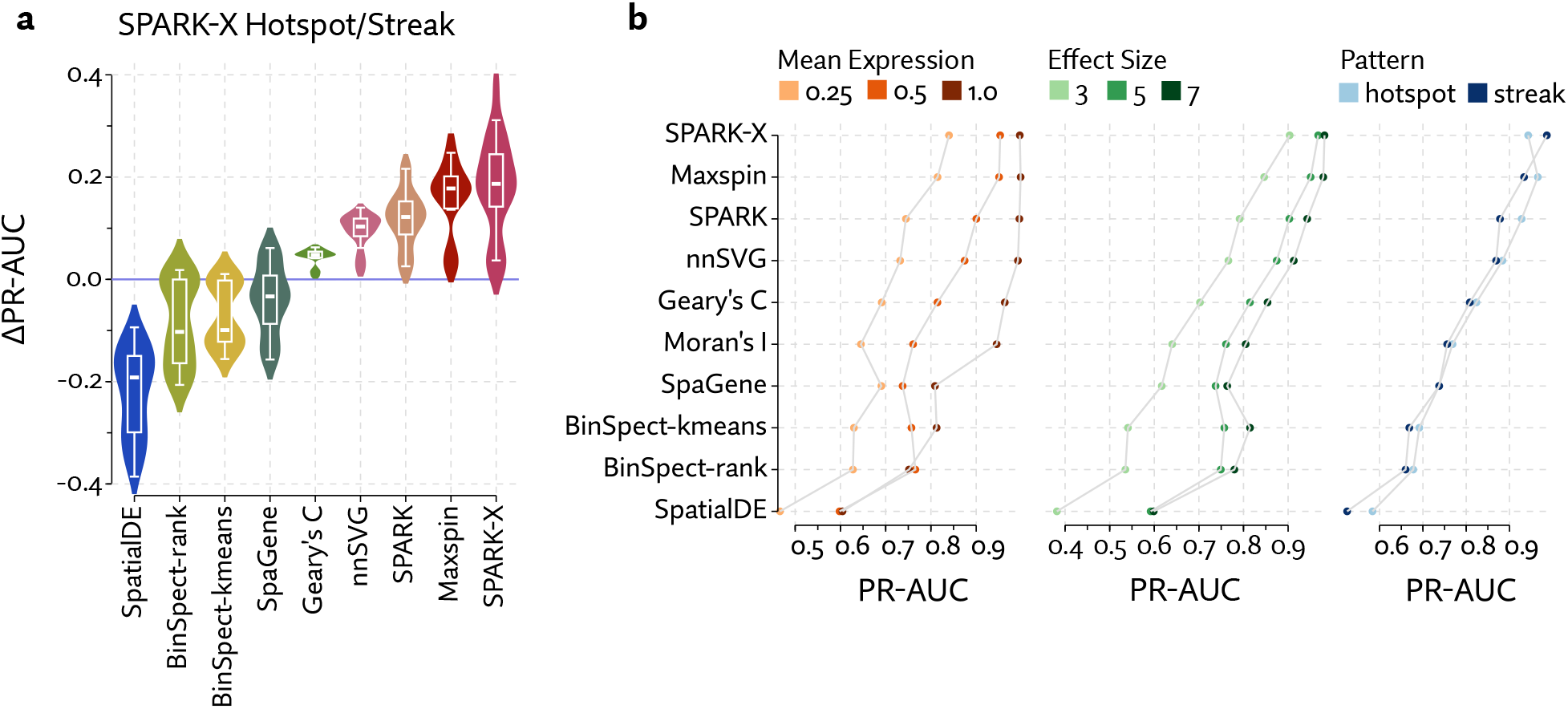
A simulation consisting uniformly distributed cells with a circle (“hotspot”) or vertical band (“streak”) of perturbed expression. **a** Aggregate performance is measured by computing the area under the precision-recall curve and subtracting the value obtained by Moran’s I. **b** Performance is further broken down by simulation parameters.

### CosMx SMI

CosMx Spatial Molecular Imager (SMI) a recently introduced platform from NanoString for spatial profiling of RNA and protein expression. We used CosMx data from He et al. [2022] to evaluate detection of spatially varying genes. Expression of 960 genes was measured across roughly 800 thousand cells, across 8 human non-small cell lung cancer samples. We relied on the authors annotation of tissue microenvironments. We tested individual fields of view for each sample, in part to create a variety of scenarios, but also to allow SPARK and SpatialDE to be run on this data, as they ordinarily would struggle to scale to this many cells. Our ground truth set of spatially varying genes consisted of any gene that varied between any pair of microenvironments, which was on average 419 per field of view.

Maxspin had consistently higher performance that competing methods (Figure 2c). SPARK-X sometimes improved on Moran’s I significantly, but as with the Visium data, it is inconsistent, producing poor results on other samples. They remaining methods show performance close to or sometimes below that of Moran’s I. As with the MERFISH data, and in stark contrast to the Visium data, SpatialDE showed extremely poor performance, with a median ΔPR-AUC of −0.197.

Figure 2d shows several examples of large disagreements between methods. Often methods will disagree on the relative significance of very small neighborhood of cells with elevated expression (e.g. CCL20), with Maxspin preferring patterns involving more cells. The Gaussian process methods seem to occasionally fail on examples with large unoccupied areas (e.g. OLFM4). SPARK-X produces some unpredictable results, failing to identify fairly obvious examples when they consist of sporadic small regions spread throughout the sample (e.g. ITGA3, TPSB2).

The probe set include 20 negative control probes, which we expect to be unexpressed. Thus, any nonzero counts for probes represents pure technical noise that should not be detected as spatially varying. Figure 2e shows that while these are typically ranked lowly by all methods, the median rank for Maxspin is the lowest, with nnSVG very nearly tied.

In Supplemental Figure 3, we show additional examples of spatial information scores computed by Maxspin, on small sections of one sample. The highest scoring genes with tumor regions, unsurprising giving its specific spatial locality.

### Simulations

To explore a variety of scenarios with a known ground truth, we simulated spatial expression data. The simulation assumes some number of discrete cell types and genes. Random expression values were drawn from negative binomial distributions. For 7000 of the 8000 simulated genes, this distribution is fixed across cell type, and for the remaining 1000 it is perturbed, representing a ground truth subset of spatially varying genes. To simulate the spatial arrangement of cells, we developed a Cellular Potts model [Graner and Glazier, 1992], in which cells migrate and deform according to an actin-inspired mechanism [Niculescu et al., 2015]. Cells adhere to one another with greater or lesser strength according to a random cell type affinity matrix.

To consider conditions of more or less spatial coherence we initialized cell positions into highly ordered arrangements, by assigning cell type according to sine waves varying across the spatial dimensions, producing homogeneous clumps of cells. Simulation parameters were then chosen so that cell arrangements slowly grew more disordered with more iterations, while still displaying some patterning due to varying cell type adhesions. Thus, running the simulation for a larger number of iterations produces a arrangement of cells with spatial patterns that are subtler but still present (Supplemental Figure 5). We ran a variety of simulations with 2000 cells and 10000 cells. Because SPARK and SpatialDE are both Gaussian process methods, the former with O(n^3^) inference, and the latter limited by GPU memory, they were unable to scale to the n = 10000 simulations.

Maxspin was found to outperform other methods by a large margin in these simulations (Figure 3a,b). These results hold up across a variety of parameterizations of the simulation (Figure 3c). SPARK and nnSVG compete for second, but SPARK fails to scale to 10000 cells, and SPARK-X shows wildly variable performance. Geary’s C slightly improves on Moran’s I, prehaps due to more subtle local variation, while all other methods consistently underperform Moran’s I.

The ground truth in the real data sets examined in previous sections was constrained to fairly obvious examples found by calling differential expression between large annotated regions. Because the ground truth is known here without any intermediating statistical procedure, it contains more subtle examples, so differences in accuracy are amplified here compared to real data, explaining the larger improvement we see with Maxspin.

We additionally adapted simulations from Zhu et al. [2021] consisting of cells distributed uniformly at random across a rectangle, with a subset of spatially varying genes that have modulated expression either in a single vertical band region (“streak”) or a circular region (“hotspot”). We generated a number of simulations by varying parameters controlling mean expression, overdispersion, and effect size. The cosine and Gaussian coordinate transformations used by SPARK-X are well tuned to detect these particular simple spatial patterns, yet Maxspin achieves nearly equivalent performance without any specialty spatial kernels. As with the Cellular Potts simulation, SPARK and nnSVG compete for second place, Geary’s C slightly surpasses Moran’s I, and all other methods underperform Moran’s I. We might expect SpatialDE to perform similarly to the other Gaussian process methods, but it once again assigns 0 p-values to many examples, erasing any meaningful ordering.

Because most biological instances of spatially coherent expression, especially at the cellular level, are so much more complex, we believe these streak and hotspot simulations are less realistic than our Cellular Potts model, yet they do offer an important alternative perspective. SPARK-X does well here, but considerably worse on the more sophisticated Cellular Potts model, with accuracy collapsing in some cases of subtle spatial organization, while Maxspin very consistently performs well across all simulations.

### Spatially Varying Expression in Renal Cell Carcinoma

We used Maxspin to investigate patterns of spatially varying gene expression in a CosMx dataset of a human renal cell carcinoma (RCC) sample, consisting of 178,410 cells and 993 genes, with an average of 222 counts per cell. The data involved both a large number of cells, which many existing methods fail to scale to, and relatively sparse counts, introducing uncertainty which many methods fail to account for. Maxspin is well suited to this data, as it scales to very large number of cells, and is able to account for uncertainty under a variety of distributional assumptions.

We first identified cell types using unsupervised clustering of expression data alone. Count data was normalized using Sanity [Breda et al., 2019], then clustered using Leiden [Traag et al., 2019]. This produced 24 clusters, which we assigned names to by comparing average expression across clusters, making use of an existing catalog of human kidney cell types [Stewart et al., 2019] (Figure 5c).

**Figure 5:**
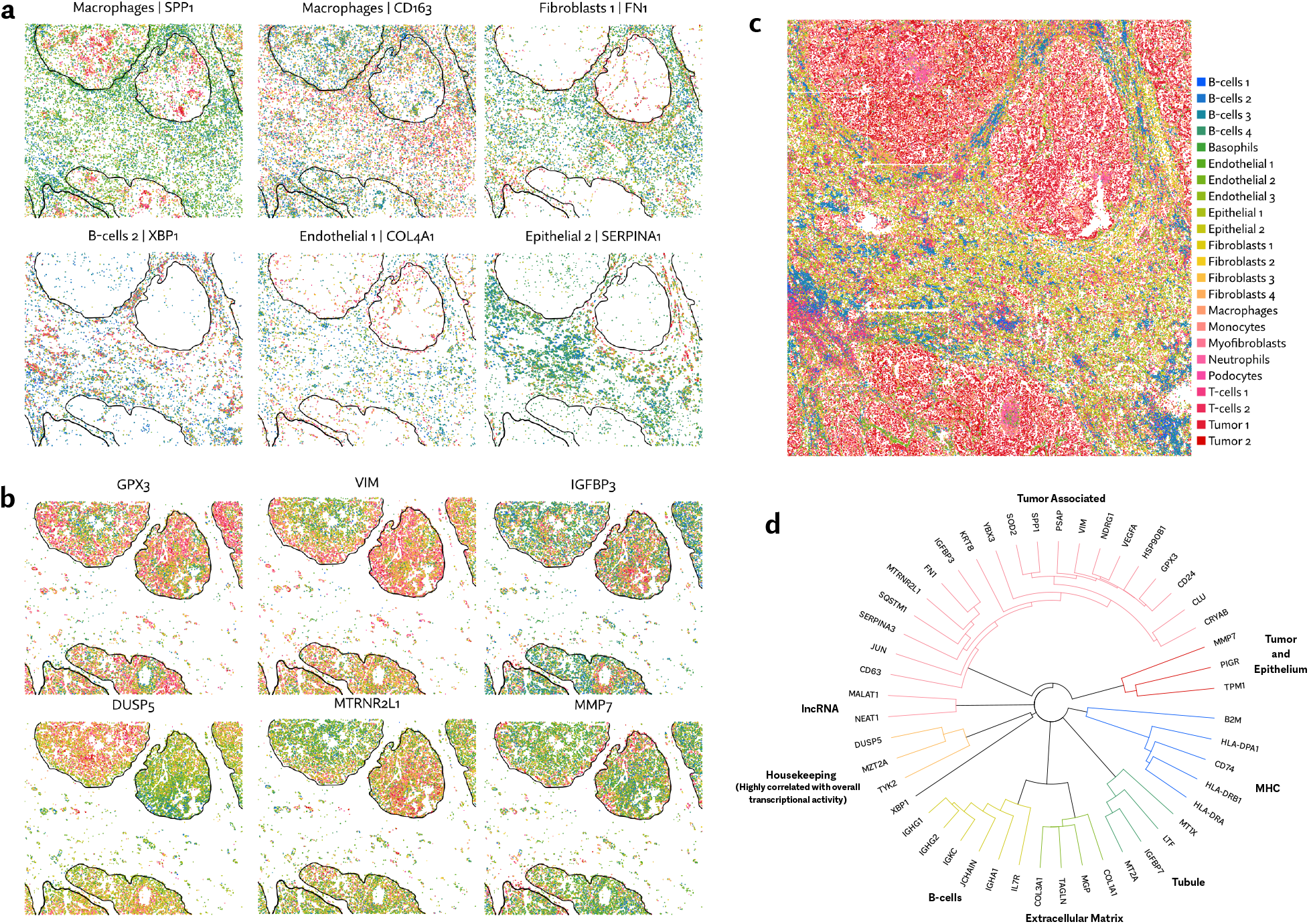
**a** Spatially varying genes detected in non-tumor cell types. Major tumor regions are indicated by outlines. **b** Spatially varying genes detected in tumor cell types. **c** Classification of cell types using a unsupervised, spatially unaware clustering. **d** Hierarchical clustering of spatially varying genes using pairwise spatial information.

Computing spatial information across all cells will often recapitulate markers for known cell types. For example, the highest scoring gene in this dataset is the immunoglobulin gene IGHG1, revealing populations of B-cells which we could have annotated by other means. To reveal more subtle spatial patterns, we instead separately ran Maxspin on each cluster to discover patterns of spatial variation that are less visible to standard clustering methods. Many examples of spatially coherent intracluster variation were found in the tumor cell clusters (Figure 5b).

Many of these genes have been studied in the context of RCC. For example, GPX3, VIM, IGFBP3 are known to be important markers in clear cell RCC [Miess et al., 2018, Williams et al., 2009, Chuang et al., 2008], but much less is known about their patterns of spatial expression within the tumor. DUSP5, which in varying contexts can act either as a tumor suppressor or promoter [Kutty et al., 2017], is seen to be highly expressed specifically in the upper-left tumor region. Curiously, the adjacent tumor region shows distinctly high expression of the MT-RNR2 like-1 pseudogene. MMP7 is upregulated along the tumor-stroma boundary, consistent with its role in degrading extracellular matrix to facilitate tumor invasion [Yokoyama et al., 2008].

In non-tumor cell clusters, a number of interesting examples of spatially varying genes also emerge (Figure 5a). Tumor associated macrophages disproportionately show upregulation of SPP1 and downregulation of CD163, consistent with prior findings [Nirschl et al., 2020, Peranzoni et al., 2018]. Along the tumor-stroma boundary, extracellular matrix genes fibronectin (FN1) and collagen type 4 alpha 1 (COL4A1) are highly expressed in fibroblasts and endothelial cells, respectively. The transcription factor X-box binding protein 1 (XBP1), which is involved in plasma cell differentiation [Reimold et al., 2001], shows spatially varying clusters of high expression within one of the B-cells clusters. Within an epithelial cell cluster, SERPINA1 shows a elevated expression surrounding one of the tumor regions, but not the others.

For several of these genes Supplemental Figure 6 shows disaggregated information scores, which can be plotted as saliency maps providing clear visual explanations for why a gene scores highly. Examples of spatially varying genes with a span of spatial information scores are shown in Supplemental Figure 2. Not surprisingly, high expression is typically a necessary condition for a high information score (Supplemental Figure 1), but not a sufficient condition, as some high expression genes have relatively low spatial information.

**Figure 6:**
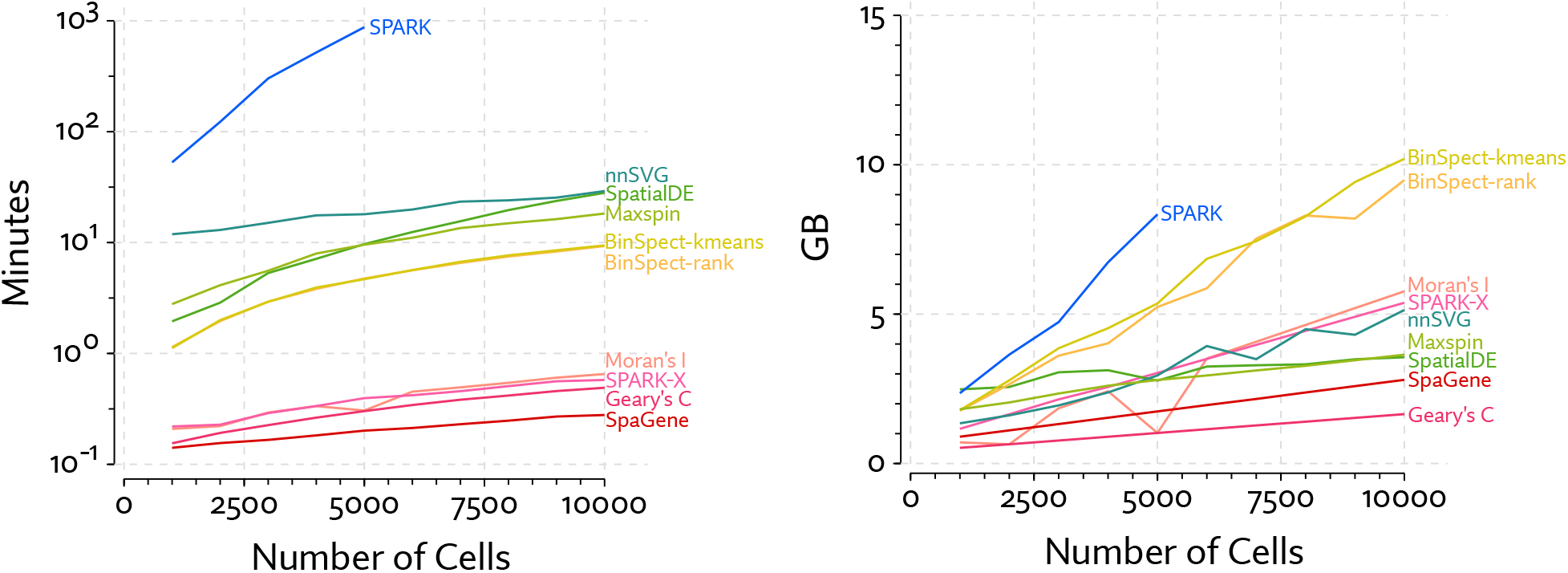
Run time and system memory usage were measured for each method across simulated data sets with varying numbers of cells. Most methods scale linearly with the number of cells, whereas SPARK and SpatialDE have observable different trajectories as they rely on Gaussian process models with exact inference.

The spatial auto-mutual information score can be trivially generalized to compute the mutual information between a pair of genes. Doing this for all O (n^2^) pairs is typically intractable unless n is small, but if we first filter to consider only genes with high auto-mutual information, this becomes a viable way to explore the spatial relationships between genes, which avoids coercing cells into distinct types or clusters, where they might not always unambiguously fit.

We chose genes with a spatial information score above 20, resulting in 48 genes, then computed pairwise information between these genes. Hierarchical clustering of this pairwise information matrix reveals specific relationships of spatial organization among these genes (Figure 5d). Twenty of the genes were broadly tumor associated, yet as indicated by the structure of the dendrogram, show varying patterns of expression within the tumor. Outside of the tumor associated group, we see some predictable groupings which we have broadly identified as B-cell/immunoglobulin, extracellular matrix, tubule, MHC, housekeeping, and lncRNA. There is significant variation within these groups, for example the MHC I gene B2M is in the same tree as four MHC II genes, but at a much greater distance. XBP1 shows a highly distinct pattern of expression, with low spatial information with any other of the genes. Taken together, this gives a rough map of the spatial organization of a subset of genes in this dataset.

### Performance

Te evaluate computational costs of the methods, we ran them on simulated data with varying numbers of cells. Since these methods test each gene individually, they all scale linearly with with the number of genes, thus we kept the number of genes fixed at 8000.

The results of this benchmark are shown in Figure 6. SPARK was by far the most computationally expensive method in terms of time and memory. We ran benchmarks for SPARK only out to 5000 cells, as it became exceedingly time consuming. Most methods scale roughly linearly with the number of cells. The exceptions are the Gaussian process methods SPARK and SpatialDE. The one other Gaussian process method, nnSVG, manages to achieve linear time inference, but was still the second slowest method. SpatialDE overtakes nnSVG at 10,000 cells, but is faster for fewer cell in part because it runs on a GPU.

To avoid giving Maxspin any opportunity for unfair advantage, we disabled convergence checks when optimizing and let it run for the maximum number of iterations. As a result, the results reported for Maxspin are strictly an upper bound on inference time. Regardless, it significantly outperforms Gaussian process methods, though it cannot match the performance of methods like Moran’s I, with simple closed form computations.

SpatialDE and Maxspin differ from the other methods in that they run on GPUs. Though it is possible to run Maxspin without a GPU, it is not designed or optimized to do so, so we did not systematically benchmark this option. In limited tests, it was roughly 32 times slower. While not advised, running Maxspin without a GPU is still considerably faster than SPARK.

GPU methods also have work within the constraints of GPU memory, which is typically much more limited than system memory (e.g. 12GB GPU memory vs 64GB system memory in the system we used for these benchmarks). Maxspin is designed to split work up and run in batches avoiding an issue with memory constraints until spatial transcriptomics technology scales to orders of magnitude larger. SpatialDE on the other hand, does rapidly run up against this limitation. Indeed, memory usage is inherently higher in Gaussian process inference, even when approximate inference methods are used.

All benchmarks were performed using Nvidia GeForce 3080 Ti GPU and AMD Ryzen 5950X CPU.

## Discussion

The spatial auto-mutual information score computed by Maxspin appears to better capture spatially coherent expression patterns than existing methods across various datasets and simulations. When presenting the results as ΔPR-AUC values, quantifying the improvement over Moran’s I, a simple and well established definition of spatial correlation, we see that specialized modern methods are not always an improvement. Methods aimed at detecting spatially varying genes are sometimes guilty of disregarding decades of work in spatial statistics and fail to demonstrate a consistent advantage.

Taking the benchmarking results together, we see some patterns emerge. Methods which rely on binarization of expression data, SpaGene and BinSpect, clearly discard some useful information and consistently underperform Moran’s I. Gaussian process methods — SpatialDE, nnSVG, and SPARK — often perform well. In the case of SpatialDE, it performed very well on the Visium benchmark, and very poorly elsewhere, producing a preponderance of 0 p-values, which is a numerical issue that could conceivably be fixed. SPARK, which uses exact inference, rapidly becomes intractable for large numbers of cells. SpatialDE also fails to scale, though it uses approximate inference. This leaves nnSVG as the clear state of the art in Gaussian process based tests. However, each Gaussian process model lags behind Maxspin. This is in part due to Maxspin fitting a discriminative model, rather than the generative models the Gaussian process methods involve. Weaker distributional assumptions are involved, making it less prone influence by small numbers of outlier cells.

SPARK-X is an appealing alternative, which is highly efficient and adept at detecting certain spatial patterns. The set of coordinate transformation kernels it uses yields a test that is highly sensitive to, for example, circular regions and bands that span the sample, as demonstrated most distinctly in their simulation, but also in the Visium and and MERFISH brain samples, which have similar bands of distinct cell types. When spatial patterns get more subtle and local, most clearly shown in the Cellular Potts simulation, it can entirely fail to detect them. Maxspin excels in this setting, and also very nearly matches SPARK-X in sensitivity to very simple, broad patterns.

Maxspin also offers improved explainability compared to more opaque statistical models. Because the information score is a sum across cells, the individual cell-level values can be visualized, showing precisely which cells and regions contribute to a high information score (Supplemental Figure 6). No other method yet proposed can be disaggregated in this way.

In our analysis of renal cell carcinoma using CosMx SMI we demonstrated how Maxspin can be used to uncover patterns of expression in a high-dimensional, yet relatively sparse dataset, by scaling linearly with the number of cells and accounting for uncertainty in expression estimates. Though we largely focused on auto-mutual information, pairwise mutual information is a simple extension that allows genes to be clustered into an atlas of similar and dissimilar spatial patterns. As demonstrated by first clustering into cell types, then computing spatial information, spatial analysis often uncovers patterns of expression that traditional clustering approaches are insensitive to. The clearest indication of this is the several examples of spatially organized expression within the tumor regions. This variation in expression resists explanation by clustering into cell types, and is revealed most clearly by gene-level spatial analysis.

Ultimately, “spatial coherence” or “spatially varying” expression can be subject to definitional disagreement. There may be different defensible answers to what constitutes a meaningful pattern of spatial organization. Other fields have grappled with these issues long before the recent relevance of spatial statistics in genomics. For example, the question of how best to measure segregation has been debated for decades in spatial demography [Wong, 2016]. The benchmarks presented here all rely on calling differential expression between identified transcriptionally distinct regions. While unlikely to capture all meaningful spatial variation, it is an intuitive baseline that we expect captures the most significant variations that should be uncontroversial as examples of SVGs. Further, these results hold up in our simulations in which the ground truth is known.

The principle presented here, using a discriminative model to maximize a bound on mutual spatial information, has other potential uses which we have yet to fully explore. An obvious extension is to directly control for possible covariates, by estimating conditional mutual information. This is straightforwardly achievable is by fitting the model twice, once including expression values and the covariate, and again with just the covariate, and subtract the two, relying on a basic identity of conditional mutual information. A less trivial extension is to use maximization of spatial mutual information as an objective function in gating, clustering, or dimensionality reduction tasks. This adds a level of complexity, because we would be simultaneously optimizing a function to assign cells to clusters (or to low dimensional representations) and the discriminative bound on on mutual information, but this is not so different than the set up in generative adversarial networks [Goodfellow et al., 2014], which have seen tremendous success, despite there being a level of trickiness training such models.

By framing spatial coherence as spatial mutual information, and developing the tools for practical inference, we have presented a promising new perspective on spatial expression analysis, and have have taken the first step exploring this idea by demonstrating a simple discriminative model that is more efficient and more accurate than state of the art Gaussian process models at identifying spatially varying genes across many settings. Though not a replacement in all cases for generative probabilistic models, it is in many cases a more accurate and efficient alternative that is designed to scale with spatial transcriptomics technology as it inevitably expands to very large data sets in the coming years.

## Online Methods

### Spatial auto-mutual information

The concept of spatial auto-correlation has been studied for decades using metrics like Moran’s I [Ripley, 2005]. Our goal is to create a more general and flexible notion spatial dependence by instead adopting an information theoretic perspective and considering auto-mutual information.

We score the “spatial coherence” of a gene’s expression by approximating the spatial auto-mutual information over nearby expression values. Let p(x) be the marginal expression distribution for the gene of interest, p_N_(x,x′) the joint distribution over pairs of expression values occurring in the same spatial neighborhood. The *spatial auto-mutual information* is then

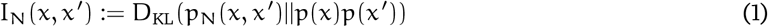

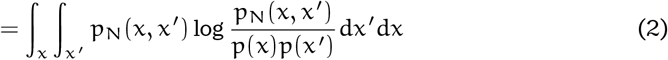

where D_KL_ is the Kullback-Leibler divergence.

This definition hinges on how “spatial neighborhood” is defined. In the simplest form, we can consider two cells/spots neighbors if they are within some distance ∊. If s, s′ ∈ ℝ^d^ are spatial coordinates corresponding to x, x′, respectively, we can marginalize out the spatial coordinates within the same neighborhood to arrive at a joint neighborhood expression distribution on fixed radius neighborhoods

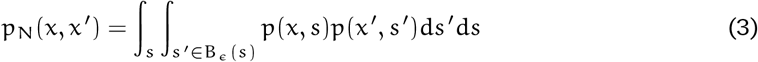

This can be generalized by defining soft neighborhoods using a joint distribution over pairs of spatial positions

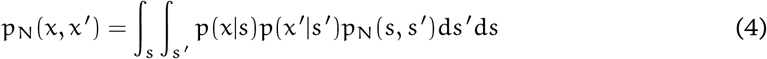

where p_N_(s, s′) is chosen to assign higher probability to nearby coordinates. To recover Equation 3, we can make it a uniform distribution over pairs within distance ∊ of each other.

A related notion of spatial mutual information occurs in Altieri et al. [2018]. There, spatial mutual information is defined simply as the mutual information between a pair of observed values (x, x′) and their spatial distance d(s, s′), both of which are discretized in their setting. This is seemingly more straightforward as it avoids having to define a neighborhood distribution, but in a large dataset training on all n^2^ pairs would be inefficient, and randomly sampled subsets of pairs will tend to be mostly distant. So in practice, this is potentially harder to estimate. Furthermore, the precise distance between a pair of cell is unlikely to be meaningful when those cells are far away from each other.

Another possible definition is to consider the mutual information between a single expression value and its entire spatial neighborhood. This may enable the detection of more elaborate spatial patterns, but has practical issues. Neighborhoods could easily be memorized by a model, overestimating mutual information, so the joint distribution would have to be very carefully modeled to avoid overfitting. We would also have to aggregate information across every node’s neighborhood, using for example a graph neural network [Kipf and Welling, 2016], at every step of training and evaluation. The computational cost would thus limit how large of a neighborhood we could consider.

Computing mutual information has two major hurdles in practical applications. First, it necessitates finding a reasonable model for the joint and marginal probabilities. An overparameterized model for p_N_(x, x′) risks memorizing the data or otherwise discovering spatial patterns that are too subtle to be meaningful, whereas an insufficiently expressive model will fail to find true patterns. Second, even with a suitable model in hand, we still have to compute the integral in Equation 2. In this work, we sidestep both of these issues by instead estimating a lower bound on the mutual information using a simple discriminative model.

### Approximating mutual information

A number of recent works have focused on avoiding the problem of computing mutual information directly by deriving more tractable lower bounds [Tschannen et al., 2019]. These formalize the intuition that the easier it is to distinguish random draws form the joint distribution from random draws from the marginals, the higher the mutual information must be. Or, in the context of spatial transcriptomics: a gene’s expression is spatially coherent if we can, with some reliability, predict whether a pair of expression values was drawn at uniform, or drawn from nearby cells.

A number of bounds on different divergence measures have been derived (for a review and comparison, see Tsai et al. [2021]). For this work, we found the bound on Jensen-Shannon divergence proposed by Nowozin et al. [2016] to work well, which is equivalent to the binary cross-entropy objective used by Brakel and Bengio [2017]. The Jensen-Shannon mutual information can be defined as the Jensen-Shannon divergence between P_N_(x, x′) and p(x)p(x′), and bounded below by

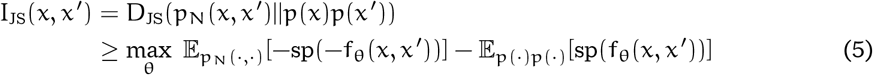

where sp(x):= log(1+exp(x)) is the softplus function, and f_θ_ is a classifier function with parameters θ. Both expectations are taken over (x,x′) with respect to either the joint neighborhood distribution or the product of the marginals.

Intuitively, we want to train f_θ_ to distinguish samples drawn from the same neighborhood, using P_N_, and pairs that are independent draws from across the sample. The better it is able to distinguish these, the higher the lower bound on mutual information. Thus, expression patterns are spatially coherent to the degree that we can learn a pattern in nearby pairs.

Assuming we can efficiently sample from p and p_N_, and f_θ_ is differentiable with respect to θ, this objective can be easily optimized using stochastic gradient descent, by drawing samples from the joint and marginals at each iteration, computing the objective in Equation 5, and backpropagating gradients to update θ.

Concretely, each step of the optimization algorithm proceeds as follows, where x = (x_1_,…, x_n_) is a single gene’s expression measured across n cells or spots, and s = (s_1_,…, s_n_) is their spatial positions.

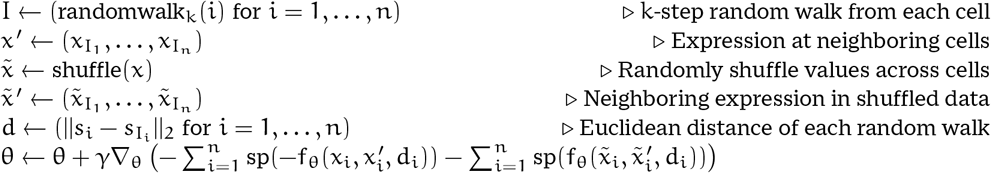

Here γ is the step size. In practice we use the Adam [Kingma and Ba, 2014] optimizer, rather than a fixed step size.

### Choosing neighborhood joint distribution

The definition of “spatial neighborhood” is determined by the neighborhood joint distribution P_N_. Optimizing the Jensen-Shannon lower bound using stochastic gradient descent necessitates a distribution that can be efficiently sampled from. In Equation 4 we showed how joint neighborhood expression can be defined in terms of a joint distribution over pairs of positions p(s, s′) defining a soft neighborhood, where s, s′ are spatial positions of cells.

In Maxspin, we use the distribution induced by considering random walks of a fixed length between s and s′.A neighborhood graph can be formed by Delaunay triangulation, fixed distance neighborhoods, or according to a grid in the case of Visium data. From this graph, a transition matrix is formed where each neighbor is visited with equal probability. We can then define the joint distribution over positions as

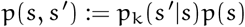

where p(s) is a uniform distribution over cell positions, and p_k_(s′|s) is the probability of starting at the cell at position s and arriving at the cell at position s′ after k random steps.

Larger values of k will consider a larger neighborhood, and thus be more tuned to finding larger scale spatial patterns, and a small k will be more tuned to finding small scale patterns. Throughout this paper we used k = 10, finding the results to be not particularly sensitive to this parameter, so long as it was not very small (less than 3). The limiting stationary distribution is a uniform distribution over pairs of nodes, so at a certain point performance will degrade if k is too large, but the computational burden of sampling such long walks becomes an issue before that point.

This random walk distribution is largely equivalent to the p-step random walk kernel discussed by Smola and Kondor [2003], which is proposed as a more practical alternative to their graph diffusion kernel.

It has yet to be established definitively whether definitions of spatial locality that are graph based, as used here, or distance based, as used for most Gaussian process covariance functions, are a better fit for spatial transcriptomics in general. Graph based approaches are simpler though, since there is no need to try to calibrate the scale of the kernel function to the spatial coordinates which come in a variety of units (pixels, grid indices, micrometers, etc). As with common distance based kernel functions, random walk based kernels also emphasize nearness, as a walk between two distant cells is less probable.

### Choosing a classifier

Bounding mutual information using a classifier solves the issue of having to model the joint probability distribution. Classifier models, typically conceptualized as a distribution over labels conditioned on observations, are typically far easier to model than full generative probabilistic models. Framing the problem as classification also presents an opportunity to remedy another major weakness of mutual information methods: since mutual information captures any non-independence relationship it can find relationships that are spurious. In principle, any differentiable classifier can be used for fθ, but too powerful a learner, like a deep neural network, risks memorizing the data, or identifying patterns too subtle or elaborate to be of interest. In practice we need a learner than is sufficiently weak, yet able to capture the kinds of patterns that are of potential scientific interest.

Since we are most often interested in the phenomenon of a gene being expressed similarly in spatial neighborhood, we use a simple classifier of the form

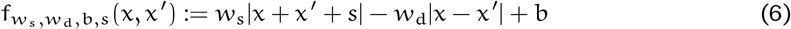

We constrain the weights w_s_, w_d_ ∈ ℝ^+^ to be positive using a softplus transformation. The intuition here is that we assign high scores to node pairs that are similarly expressed (captured by the w_d_|x − x′|term) and both either unusually low or high (captured by the w_s_|x + x′ + s| term). Dropping the positivity constraint would allow the detection of anti-correlation, which could potentially be of interest, but more often we are interested in positive correlation, and restricting the signs of the weight slightly improves performance for such cases.

We found this function family to be suitably general to discover simple patterns of low or high expression regions, but in other settings where the goal is to find more elaborate spatial patterns, the method can be trivially adapted by choosing a more powerful classifier. The risk of doing so is to overfit or memorize the data, but this can be further remedied with standard regularization techniques.

We further allow the classifier weight these scores according to the distance of the random walk, by passing the distance of the random walk to classifier for both the neighborhood and uniform sampled pair. This allows it to discount long walks when the spatial organization is very local. To do this we weight classifier scores by exp(−d/α), where d is the Euclidean distance of the walk, and α is a parameter that is learned on a per-gene basis. The weighted classifier function then takes the form

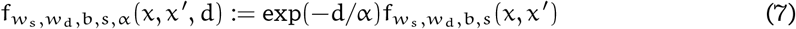

Optimization is over the set of parameters θ = (w_s_, w_d_, b, s, α), each a real number.

### Accounting for uncertainty in count data

Discrete and often sparse count data represents a noisy estimate of gene expression. Methods often fail to account for this source of uncertainty resulting overconfident predictions. Instead of training on raw, or transformed count data, we instead resample expression values periodically during stochastic gradient descent, effectively optimizing an expectation over unobserved expression values. This has the concrete effect of minimizing the possibility of overfitting.

With count data, we can either assume counts represent an unconfounded measurement of absolute expression, or we can assume that the total counts per cell/spot represent a confounding variable and instead deal in proportional expression (e.g. “transcripts per million”). In the former case, we adopt a Gamma-Poisson model, and train on values sampled from the posterior Gamma distribution at each iteration. When treating data proportionally, we adopt a Dirichlet-Multinomial model and train on samples drawn from the posterior Dirichlet distribution.

The same approach can be used with a more sophisticated probabilistic model, by running Maxspin on samples drawn from its posterior. We took this approach in our analysis of the CosMx RCC data. Posterior inference done by Sanity [Breda et al., 2019] provides posterior means and standard deviations. We used these values to sample expression values from a normal distribution when running Maxspin. In the future, other types of models can be used to handle isoform- or allele-specific expression by accounting for uncertainty over read assignment.

### Improving performance in sparse data by binning

When expression for a gene is very sparse, training can be inefficient, because it relies on the relatively small proportion of random walks between cells with nonzero expression. To remedy this, we bin data at multiple resolutions, and train jointly on the original data and the binned data.

Binning is nontrivial because if bins end up with differing numbers of cells, it can introduce subtle statistical dependencies which can falsely manifest as positive spatial information scores, even if normalizing for cell count. To overcome this, we spatially bin data using a modified kd-tree. For n cells/spots and a specified bin size k, we first discard a random n mod k cells to ensure the data can be exactly binned. We then recursively split along alternating axes, ensuring at each split that both partitions can be divided exactly by k, until we reach partitions of size k. The result is spatially arranged bins each with exactly k cells. By default we include binned data with bin sizes k = 4, 8, 16 when training. Of course, the result of binning is fewer bins than there were cells, so when training upweight the objective function for each binning by 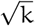. Weighting offers a trade-off. More highly weighted large bins will make the method more sensitive to broad but sparse patterns, but somewhat less sensitive to very small scale patterns.

### Hierarchical Clustering with Pairwise Spatial Information

Pairwise information scores were computed and hierarchical clustering was run with with complete linkage. Pairwise information, unlike distance larger for more closely related genes. To use with hierarchical clustering, which assumes pairwise distances, pairwise information I_ij_ is first symmetrized as 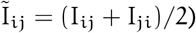 to reduce numerical inaccuracy, then transformed into a distance as

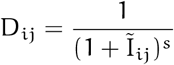

where s is a constant controlling the relative scaling, which we set to s = 0.5.

### Implementation

Maxspin is available as a Python package at https://github.com/dcjones/maxspin. The Cellular Potts model used is simulations is available as Julia package at https://github.com/dcjones/Cronenberg.jl.

The method in implemented in Python using jax [Bradbury et al., 2018] and flax [Heek et al., 2020], and built on data structures from squidpy [Palla et al., 2022].

## Acknowledgements

This research was supported by funding from the National Institutes of Health (P01-CA225517 and U19AI128914, D.J., R.G., E.W.N.), the Immunotherapy and Translational Data Science Integrated Research Centers at Fred Hutchinson to D.J., R.G., E.W.N., the Fred Hutchinson Cancer Center New Development funds (E.W.N.), and The Andy Hill Endowment Distinguished Researcher CARE fund (E.W.N.). R.G.’s research was supported by institutional funding from the Faculté de Biologie et Medicine of the University of Lausanne, and the Lausanne University Hospital. We are grateful to Edward Zhao for helpful feedback on the manuscript.

## References

L Altieri, D Cocchi, and G Roli. A new approach to spatial entropy measures. Environ. Ecol. Stat., 25(1):95–110, March 2018.

Alma Anderson and Joakim Lundeberg. sepal: Identifying transcript profiles with spatial patterns by diffusion-based modeling. Bioinformatics, March 2021.

Mohamed Ishmael Belghazi, Aristide Baratin, Sai Rajeswar, Sherjil Ozair, Yoshua Bengio, Aaron Courville, and R Devon Hjelm. MINE: Mutual information neural estimation. January 2018.

Nuha BinTayyash, Sokratia Georgaka, S T John, Sumon Ahmed, Alexis Boukouvalas, James Hensman, and Magnus Rattray. Non-parametric modelling of temporal and spatial counts data from RNA-seq experiments. Bioinformatics, 37(21):3788–3795, July 2021.

James Bradbury, Roy Frostig, Peter Hawkins, Matthew James Johnson, Chris Leary, Dougal Maclaurin, George Necula, Adam Paszke, Jake VanderPlas, Skye Wanderman-Milne, and Qiao Zhang. JAX: composable transformations of Python+NumPy programs, 2018. URL http://github.com/google/jax.

Philemon Brakel and Yoshua Bengio. Learning independent features with adversarial nets for nonlinear ICA. October 2017.

Jérémie Breda, Mihaela Zavolan, and Erik van Nimwegen. Bayesian inference of the gene expression states of single cells from scRNA-seq data. December 2019.

Shang-Tian Chuang, Kurt T Patton, Kristian T Schafernak, Veronica Papavero, Fan Lin, Robert C Baxter, Bin Tean Teh, and Ximing J Yang. Over expression of insulin-like growth factor binding protein 3 in clear cell renal cell carcinoma. J. Urol., 179(2):445–449, February 2008.

Ruben Dries, Qian Zhu, Rui Dong, Chee-Huat Linus Eng, Huipeng Li, Kan Liu, Yuntian Fu, Tianxiao Zhao, Arpan Sarkar, Feng Bao, Rani E George, Nico Pierson, Long Cai, and Guo-Cheng Yuan. Giotto: a toolbox for integrative analysis and visualization of spatial expression data. Genome Biol., 22(1): 78, March 2021.

Daniel Edsgärd, Per Johnsson, and Rickard Sandberg. Identification of spatial expression trends in single-cell gene expression data. Nat. Methods, 15(5):339–342, May 2018.

Goodfellow, Pouget-Abadie, and others. Generative adversarial nets. Adv. Neural Inf. Process. Syst., 2014.

F Graner and J A Glazier. Simulation of biological cell sorting using a two-dimensional extended potts model. Phys. Rev. Lett., 69(13):2013–2016, September 1992.

Shanshan He, Ruchir Bhatt, Carl Brown, Emily A Brown, Derek L Buhr, Kan Chantranuvatana, Patrick Danaher, Dwayne Dunaway, Ryan G Garrison, Gary Geiss, Mark T Gregory, Margaret L Hoang, Rustem Khafizov, Emily E Killingbeck, Dae Kim, Tae Kyung Kim, Youngmi Kim, Andrew Klock, Mithra Korukonda, Alecksandr Kutchma, Erica Lee, Zachary R Lewis, Yan Liang, Jeffrey S Nelson, Giang T Ong, Evan P Perillo, Joseph C Phan, Tien Phan-Everson, Erin Piazza, Tushar Rane, Zachary Reitz, Michael Rhodes, Alyssa Rosenbloom, David Ross, Hiromi Sato, Aster W Wardhani, Corey A Williams-Wietzikoski, Lidan Wu, and Joseph M Beechem. High-plex multiomic analysis in FFPE at subcellular level by spatial molecular imaging. January 2022.

Jonathan Heek, Anselm Levskaya, Avital Oliver, Marvin Ritter, Bertrand Rondepierre, Andreas Steiner, and Marc van Zee. Flax: A neural network library and ecosystem for JAX, 2020. URL http://github.com/google/flax.

D P Kingma and J Ba. Adam: A method for stochastic optimization. arXiv preprint arXiv:1412.6980, 2014.

Thomas N Kipf and Max Welling. Semi-Supervised classification with graph convolutional networks. September 2016.

Raman G Kutty, Marat R Talipov, Robert D Bongard, Rachel A Jones Lipinski, Noreena L Sweeney, Daniel S Sem, Rajendra Rathore, and Ramani Ramchandran. Dual specificity phosphatase 5-substrate interaction: A mechanistic perspective. Compr. Physiol., 7(4):1449–1461, September 2017.

Qi Liu, Chih-Yuan Hsu, and Yu Shyr. Scalable and model-free detection of spatial patterns and colocalization. Genome Res., 32(9):1736–1745, September 2022.

Emma Lundberg and Georg H H Borner. Spatial proteomics: a powerful discovery tool for cell biology. Nat. Rev. Mol. Cell Biol., 20(5):285–302, May 2019.

Vivien Marx. Method of the year: spatially resolved transcriptomics. Nat. Methods, 18(1):9–14, January 2021.

Kristen R Maynard, Leonardo Collado-Torres, Lukas M Weber, Cedric Uytingco, Brianna K Barry, Stephen R Williams, Joseph L Catallini, 2nd, Matthew N Tran, Zachary Besich, Madhavi Tippani, Jennifer Chew, Yifeng Yin, Joel E Kleinman, Thomas M Hyde, Nikhil Rao, Stephanie C Hicks, Keri Martinowich, and Andrew E Jaffe. Transcriptome-scale spatial gene expression in the human dorsolateral prefrontal cortex. Nat. Neurosci., 24(3):425–436, March 2021.

Heike Miess, Beatrice Dankworth, Arvin M Gouw, Mathias Rosenfeldt, Werner Schmitz, Ming Jiang, Becky Saunders, Michael Howell, Julian Downward, Dean W Felsher, Barrie Peck, and Almut Schulze. The glutathione redox system is essential to prevent ferroptosis caused by impaired lipid metabolism in clear cell renal cell carcinoma. Oncogene, 37(40):5435–5450, October 2018.

Brendan F Miller, Dhananjay Bambah-Mukku, Catherine Dulac, Xiaowei Zhuang, and Jean Fan. Characterizing spatial gene expression heterogeneity in spatially resolved single-cell transcriptomic data with nonuniform cellular densities. Genome Res., 31(10):1843–1855, October 2021.

Lambda Moses and Lior Pachter. Museum of spatial transcriptomics. Nat. Methods, pages 1–13, March 2022.

Ioana Niculescu, Johannes Textor, and Rob J de Boer. Crawling and gliding: A computational model for Shape-Driven cell migration. PLoS Comput. Biol., 11(10):e1004280, October 2015.

Thomas R Nirschl, Margueritta El Asmar, Wesley W Ludwig, Sudipto Ganguly, Michael A Gorin, Michael H Johnson, Phillip M Pierorazio, Charles G Drake, Mohamad E Allaf, and Jelani C Zarif. Transcriptional profiling of tumor associated macrophages in human renal cell carcinoma reveals significant heterogeneity and opportunity for immunomodulation. Am J Clin Exp Urol, 8(1):48– 58, February 2020.

Sebastian Nowozin, Botond Cseke, and Ryota Tomioka. F-GAN: Training generative neural samplers using variational divergence minimization. Adv. Neural Inf. Process. Syst., 29, 2016.

Giovanni Palla, Hannah Spitzer, Michal Klein, David Fischer, Anna Christina Schaar, Louis Benedikt Kuemmerle, Sergei Rybakov, Ignacio L Ibarra, Olle Holmberg, Isaac Virshup, Mohammad Lotfollahi, Sabrina Richter, and Fabian J Theis. Squidpy: a scalable framework for spatial omics analysis. Nat. Methods, 19(2):171–178, February 2022.

Elisa Peranzoni, Jean Lemoine, Lene Vimeux, Vincent Feuillet, Sarah Barrin, Chahrazade Kantari-Mimoun, Nadège Bercovici, Marion Guérin, Jérôme Biton, Hanane Ouakrim, Fabienne Régnier, Audrey Lupo, Marco Alifano, Diane Damotte, and Emmanuel Donnadieu. Macrophages impede CD8 T cells from reaching tumor cells and limit the efficacy of anti–PD-1 treatment. Proceedings of the National Academy of Sciences, 115(17):E4041–E4050, 2018.

A M Reimold, N N Iwakoshi, J Manis, P Vallabhajosyula, E Szomolanyi-Tsuda, E M Gravallese, D Friend, M J Grusby, F Alt, and L H Glimcher. Plasma cell differentiation requires the transcription factor XBP-1. Nature, 412(6844):300–307, July 2001.

Brian D Ripley. Spatial Statistics. John Wiley & Sons, February 2005.

Alexander J Smola and Risi Kondor. Kernels and regularization on graphs. In Learning Theory and Kernel Machines, pages 144–158. Springer Berlin Heidelberg, 2003.

Benjamin J Stewart, John R Ferdinand, Matthew D Young, Thomas J Mitchell, Kevin W Loudon, Alexandra M Riding, Nathan Richoz, Gordon L Frazer, Joy U L Staniforth, Felipe A Vieira Braga, Rachel A Botting, Dorin-Mirel Popescu, Roser Vento-Tormo, Emily Stephenson, Alex Cagan, Sarah J Farndon, Krzysztof Polanski, Mirjana Efremova, Kile Green, Martin Del Castillo Velasco-Herrera, Charlotte Guzzo, Grace Collord, Lira Mamanova, Tevita Aho, James N Armitage, Antony C P Riddick, Imran Mushtaq, Stephen Farrell, Dyanne Rampling, James Nicholson, Andrew Filby, Johanna Burge, Steven Lisgo, Susan Lindsay, Marc Bajenoff, Anne Y Warren, Grant D Stewart, Neil Sebire, Nicholas Coleman, Muzlifah Haniffa, Sarah A Teichmann, Sam Behjati, and Menna R Clatworthy. Spatiotemporal immune zonation of the human kidney. Science, 365(6460):1461–1466, September 2019.

Shiquan Sun, Jiaqiang Zhu, and Xiang Zhou. Statistical analysis of spatial expression patterns for spatially resolved transcriptomic studies. Nat. Methods, 17(2):193–200, February 2020.

Valentine Svensson, Sarah A Teichmann, and Oliver Stegle. SpatialDE: identification of spatially variable genes. Nat. Methods, 15(5):343–346, May 2018.

V A Traag, L Waltman, and N J van Eck. From louvain to leiden: guaranteeing well-connected communities. Sci. Rep., 9(1):5233, March 2019.

Yao-Hung Hubert Tsai, Martin Q Ma, Muqiao Yang, Han Zhao, Louis-Philippe Morency, and Ruslan Salakhutdinov. Self-supervised representation learning with relative predictive coding. March 2021.

Michael Tschannen, Josip Djolonga, Paul K Rubenstein, Sylvain Gelly, and Mario Lucic. On mutual information maximization for representation learning. July 2019.

Lukas M Weber, Arkajyoti Saha, Abhirup Datta, Kasper D Hansen, and Stephanie C Hicks. nnSVG: scalable identification of spatially variable genes using nearest-neighbor gaussian processes. May 2022.

Ariel A Williams, John P T Higgins, Hongjuan Zhao, Börje Ljunberg, and James D Brooks. CD 9 and vimentin distinguish clear cell from chromophobe renal cell carcinoma. BMC Clin. Pathol., 9:9, November 2009.

David W Wong. From aspatial to spatial, from global to local and individual: Are we on the right track to spatialize segregation measures? In Frank M Howell, Jeremy R Porter, and Stephen A Matthews, editors, Recapturing Space: New Middle-Range Theory in Spatial Demography, pages 77–98. Springer International Publishing, Cham, 2016.

Yuko Yokoyama, Frank Grünebach, Susanne M Schmidt, Annkristin Heine, Maik Häntschel, Stefan Stevanovic, Hans-Georg Rammensee, and Peter Brossart. Matrilysin (MMP-7) is a novel broadly expressed tumor antigen recognized by antigen-specific T cells. Clin. Cancer Res., 14(17):5503– 5511, September 2008.

Meng Zhang, Stephen W Eichhorn, Brian Zingg, Zizhen Yao, Kaelan Cotter, Hongkui Zeng, Hongwei Dong, and Xiaowei Zhuang. Spatially resolved cell atlas of the mouse primary motor cortex by MERFISH. Nature, 598(7879):137–143, October 2021.

Tongtong Zhao, Zachary D Chiang, Julia W Morriss, Lindsay M LaFave, Evan M Murray, Isabella Del Priore, Kevin Meli, Caleb A Lareau, Naeem M Nadaf, Jilong Li, Andrew S Earl, Evan Z Macosko, Tyler Jacks, Jason D Buenrostro, and Fei Chen. Spatial genomics enables multi-modal study of clonal heterogeneity in tissues. Nature, 601(7891):85–91, January 2022.

Jiaqiang Zhu, Shiquan Sun, and Xiang Zhou. SPARK-X: non-parametric modeling enables scalable and robust detection of spatial expression patterns for large spatial transcriptomic studies. Genome Biol., 22(1):184, June 2021.

