## Supplemental Materials for "An information theoretic approach to detecting spatially varying genes"

### Supplement
*for*
### An information theoretic approach to detecting spatially varying genes

Daniel C. Jones, Patrick Danaher, Youngmi Kim, Raphael Gottardo, Evan W. Newell

November 1, 2022

#### Additional CosMx RCC Analysis

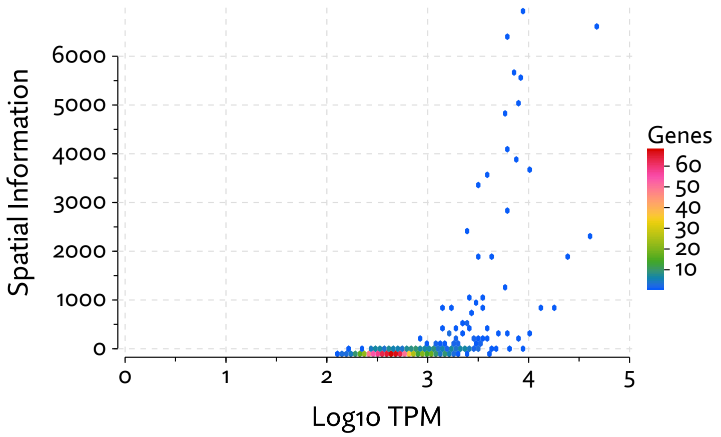


Figure 1: Spatial information scores are plotted as a function of mean expression.

#### Additional examples of spatially varying genes

Further examples of spatially varying genes and their associated spatial information scores are shown in Figures 3 and 4.

#### Interpreting the spatial information score

The spatial information score computed by Maxspin is a value $s_i \in [0, 1]$ for each gene $i$ quantifying the accuracy of predicting whether a pair of cells was sampled from a random walk in its neighborhood or uniformly at random. Though its interpretation may not be immediately obvious, it has some advantages over correlations or p-values computed by other methods.

Most critically, unlike every other method considered, the score is computed on a per-cell basis. The score for a gene $i$ is simply the sum of the scores for each cell j. That is, $s_i = \sum_j s_{ij}$. Because of this, it is trivial to determine why a gene was assigned a high score, whereas with every other existing method, some deeper analysis has to be performed. In some instances, taking the mean might be more appropriate, but by default we do not try to control for the number of cells. A spatially coherent pattern involving more cells will score higher than a similar pattern involving fewer.

These per-cell scores do have a simple interpretation as the log-probability assigned to the true labels in the classified pairs, normalized to a baseline of a random guessing. To see this we simply have to rewrite the objective function. First consider the objective function

$$s_{ij} = \mathbb{E}_{(x_n, x'_n),(x_f, x'_f)}[-\text{sp}(-f_\theta(x_n, x'_n)) - \text{sp}(f_\theta(x_f, x'_f))]$$

where sp is the softplus function, and the expectation is taken over the random walk distribution $(x_n, x'_n) \sim p_N(\cdot, \cdot)$ as well as the uniform sampling distribution $(x_f, x'_f) \sim p(\cdot)p(\cdot)$, with the mnemonic $n$ for near and f for far. This can be rewritten in terms of the sigmoid function $\sigma(x) = 1/(1 + \exp(-x))$.

$$\begin{aligned} s_{ij} &= \mathbb{E}_{(x_n, x'_n),(x_f, x'_f)} \left[\log\left(\frac{1}{1 + \exp(-f_\theta(x_n, x'_n))}\right) + \log\left(\frac{1}{1 + \exp(f_\theta(x_f, x'_f))}\right)\right] \\ &= \mathbb{E}_{(x_n, x'_n),(x_f, x'_f)} \left[\log \sigma(f_\theta(x_n, x'_n)) + \log \sigma(-f_\theta(x_f, x'_f))\right] \\ &= \mathbb{E}_{(x_n, x'_n),(x_f, x'_f)} \left[\log \left(\sigma(f_\theta(x_n, x'_n))(1 - \sigma(f_\theta(x_f, x'_f)))\right)\right] \end{aligned}$$

If we interpret $\sigma(f_\theta(\cdot))$ as the probability assigned by the classifier, we can see that this spatial information score is simply the log probability of assigning true labels to both the near and far examples. Each cell's score is then a score in $(-\infty, 0.0]$.

Random guessing would result in a baseline score of $\log(0.5 \cdot 0.5) \approx -1.38$, so scores in practice are in $[\log(0.25), 0]$, only occasionally lying slightly below due to numerical imprecision. Maxspin further normalizes these by shifting and

and scaling these to lie in $[0, 1]$, allowing them to be loosely interpreted similarly to correlations, though typically much smaller, since the classifier rarely approaches perfect accuracy. Summing across cells we then have per-gene scores $s_i$ than lie in $[0, n]$, and can be divided by $n$ if so desired.

#### Modeling concerns with count data

In the benchmarks presented we have also only considered expression variability on an absolute scale, finding genes that have consistently higher or lower counts in certain regions. This is not always the most appropriate analysis. As in other RNA-Seq protocols, in Visium what is measured is proportional, not absolute expression. The complications that come from this compositional nature have been explored in the context of traditional RNA-Seq [Quinn et al., 2018, Egozcue et al., 2020], but the situation is exacerbated in Visium because the total number of reads per-spot is at least in part a function of the degree of permeabilization of the tissue at that location, which varies by tissue type. Many methods developed have so far ignore this issue, which calls into question whether the spatial expression patterns they discover are driven entirely by gene expression or if they are being influenced indirectly by other local properties of the tissue.

In Maxspin we have implemented the option to operate on proportions, by sampling during training using a Multinomial-Dirichlet model, and adjusting by inferred or given scale factors. Another approach, which we used when analyzing the RCC data is to use a probabilistic normalization scheme that provides posterior uncertainty estimates. In general, this is our preferred approach.

A potentially more desirable alternative would be to search for spatially varying ratios of expression values in a scheme similar to what Quinn et al. [2021] proposes as an alternative to differential expression, but this will necessitate further work to overcome the obvious computational hurdles. Though we believe a compositional analysis is a more prudent approach for this type of data, we focused here on testing for patterns in absolute counts in order to create a fair comparison. Methods that normalize by total read count by default we set to us a constant size factor (rather than total number of reads) to create a fair comparison.

#### Simulation of cell positions and morphology

Our Cellular Potts model simulates cell position and morphology, with cell migrating for some number of iterations. More iterations will tend to produce less simplistic spatial patterns, white retaining a non-random configuration due to random pairwise cell type adherence. Supplemental Figure 5 shows an example of the same simulation being run increasing numbers of iterations.

#### Saliency Maps

Our spatial information score, which captures the aggregate classifier performance, can be disaggregated to cell-level scores which can be used to determine which regions are spatial coherent. The resulting images are analogous to saliency maps proposed by Simonyan et al. [2013] to visualize which regions of an image contribute disproportionately in convolutional neural network classifiers. Both here and in that work, saliency maps provide an explanation to go along with the prediction, which can be useful for diagnosis, or in the future, perhaps for clustering and other spatial analysis tasks.

Figure 6 shown examples of spatially varying genes detected in the CosMx RCC data set and their accompanying saliency maps.

#### References

Juan José Egozcue, Jan Graffelman, M Isabel Ortego, and Vera Pawlowsky-Glahn. Some thoughts on counts in sequencing studies. *NAR Genom Bioinform*, 2 (4), November 2020.

Thomas P Quinn, Ionas Erb, Mark F Richardson, and Tamsyn M Crowley. Understanding sequencing data as compositions: an outlook and review. *Bioinformatics*, 34(16):2870–2878, August 2018.

Thomas P Quinn, Elliott Gordon-Rodriguez, and Ionas Erb. A critique of differential abundance analysis, and advocacy for an alternative. April 2021.

Karen Simonyan, Andrea Vedaldi, and Andrew Zisserman. Deep inside convolutional networks: Visualising image classification models and saliency maps. December 2013.

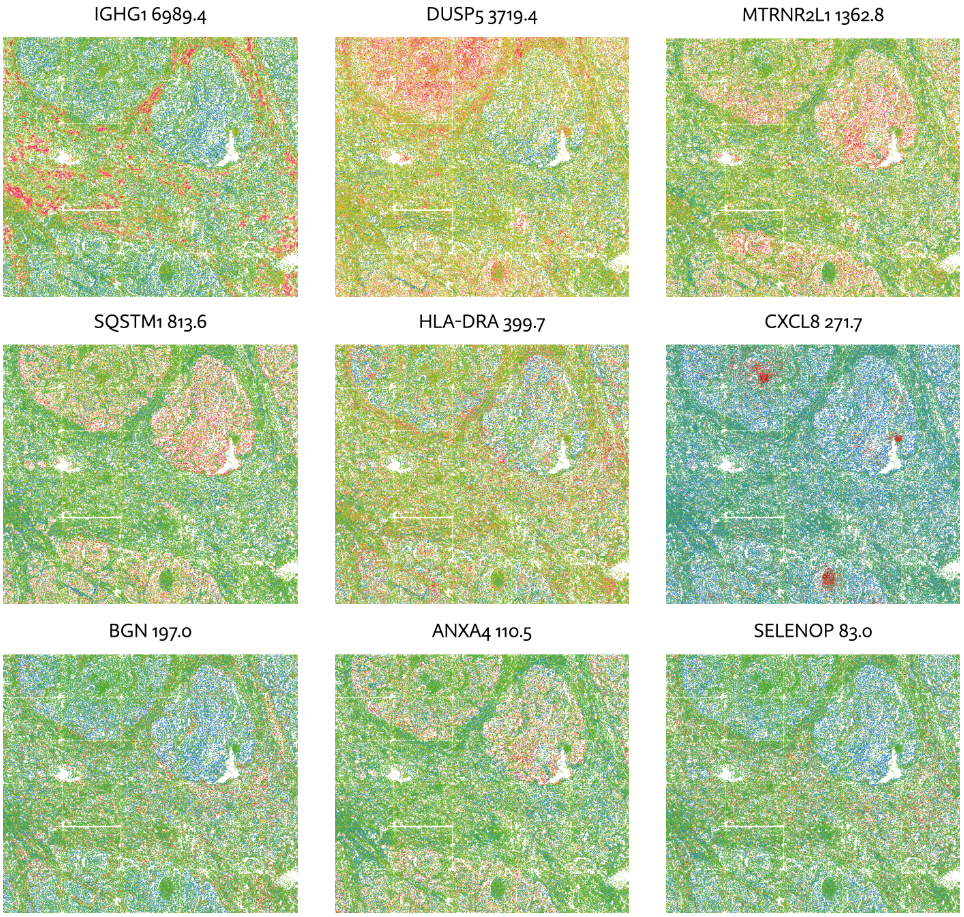


Figure 2: Examples genes with corresponding spatial information scores from the renal cell carcinoma CosMx data. These were selected to show a range of spatial information scores.

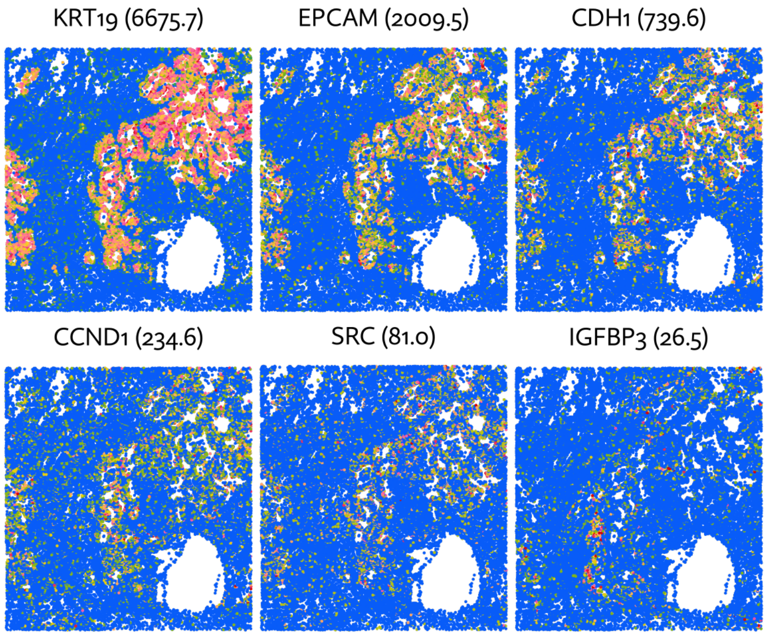


Figure 3: Examples of spatially varying gene expression along with their spatial information scores, which are seen to strongly correspond to the effect size and the number of cells involved. The highest scoring genes here show elevated expression in the tumor interior.

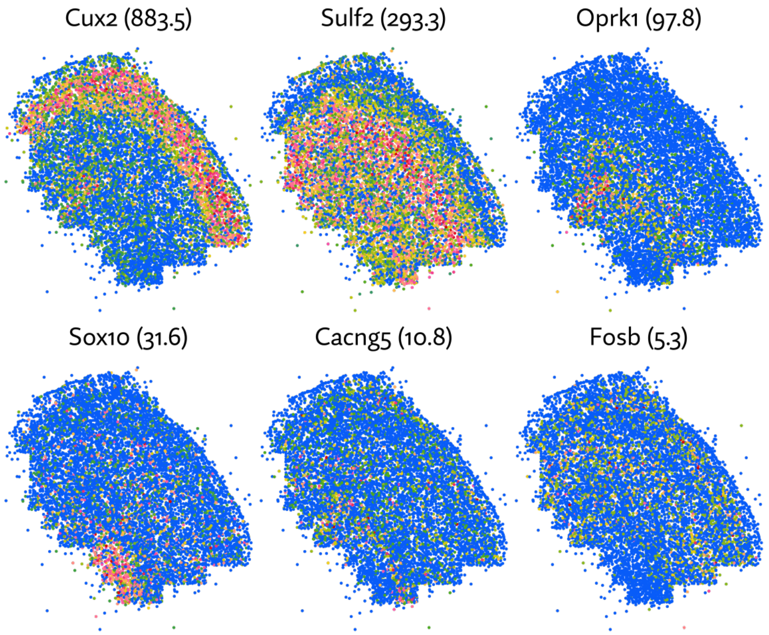


Figure 4: Examples of a range of spatial information scores assigned by Maxspin to genes in the MERFISH data set.

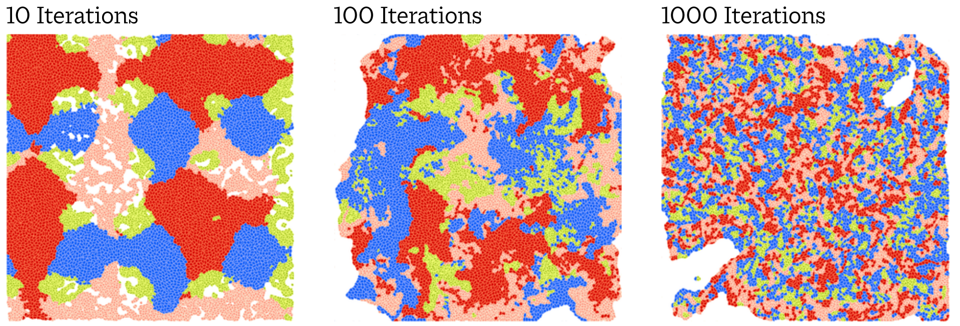


Figure 5: Cell positions and morphology are simulated using a Cellular Potts model. Cells are initially arranged by type (here indicated by color) into highly ordered "blobs". Letting the cells migrate for an increasing number of iterations produces arrangements that are less globally ordered but still governed by a pairwise cell type adhesion matrix.

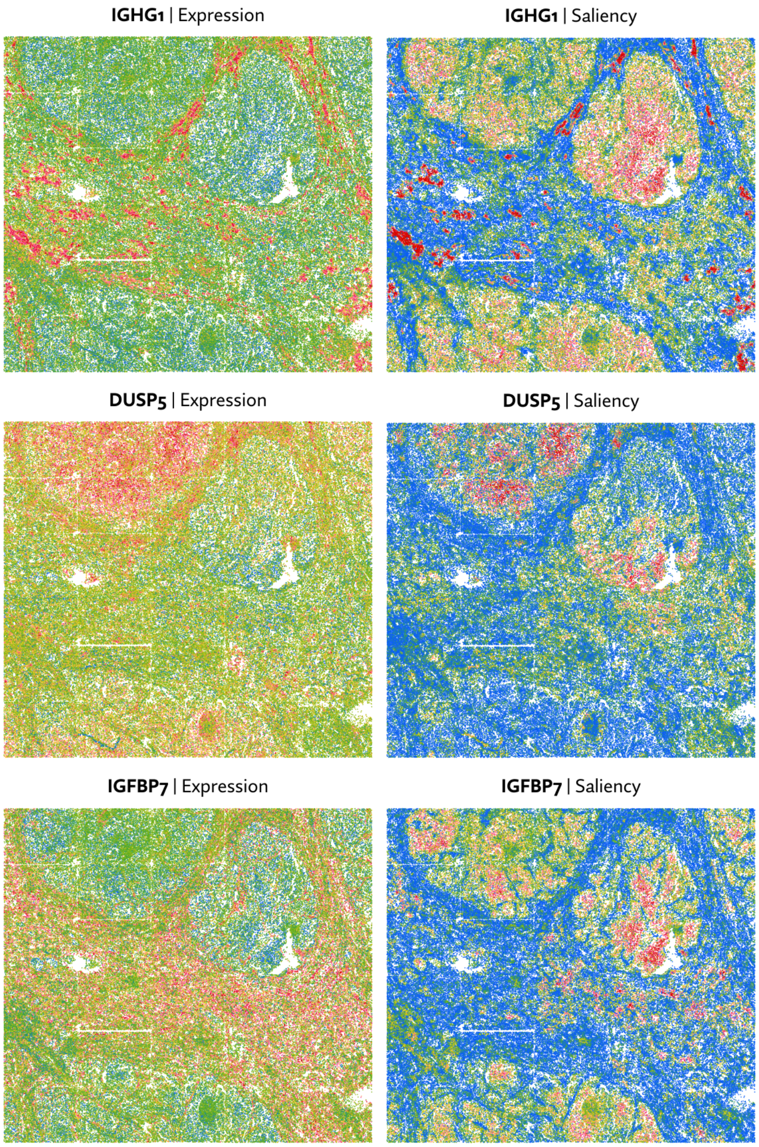


Figure 6: Expression plotted alongside saliency maps, which indicate the cell-level classifier accuracy. High saliency corresponds to regions which contribute most to the overall spatial information score.
